# Complexity shapes uniqueness: Neuropil volumes and synaptic clusters shape behavioural plasticity under challenging environments in the invasive Argentine ants

**DOI:** 10.1101/2025.03.10.642356

**Authors:** Srikrishna Narasimhan, María Eugenia Villar, Violette Chiara, Ignacio Arganda-Carreras, Sara Arganda, Magdalena Witek, Iago Sanmartín-Villar

## Abstract

Adaptation to new environments is key for organisms’ survival, but also for their invasiveness in their introduced areas. Behaviour is considered the fastest phenotype allowing adaptation, but its plasticity can involve costs as neural development. Although individuals’ investment in cognition was pointed out as unnecessary for colony behaviour in eusocial insects, recent studies are highlighting the behavioural dependence on neural traits in eusocial insects. The costs of producing behavioural and neural plastic offspring could exceed the investments of eusocial insects, in which one or certain reproductives must produce multiple offspring. Thus, we wanted to analyse the link between the neuroanatomy and the behavioural variability of *Linepithema humile*, an invasive species organised in supercolony units containing millions of individuals, to understand its adaptive mechanisms. We repeatedly tested same aged callow workers of *L. humile* in behavioural tests of increasing environmental complexity and analysed the volume of their brain functional areas (neuropils) and the synaptic clusters abundance in the mushroom body calices (information processing). Given the potential large cost of plasticity, we expected to find homogeneous interindividual neuronal structures and behavioural responses. Although *L. humile* is considered a monomorphic species, body size conditioned behavioural and neural traits and determined individuals efficiency in exploring simple environments. Contrary to our expectations, the increase in environmental complexity revealed the behavioural plasticity of *Linepithema humile* workers as well as its correlation with neuropil volumes and synaptic clusters. Our results highlight the relevance of the central complex and the mushroom bodies on exploration efficiency rather than optic and olfactory lobes. Behavioural plasticity under complex environments relied on the synaptic connections of the olfactory processing area (dense lip), while individuals with higher number of synaptic connections on the visual processing area (collar) explored complex environments less efficiently. Our results suggest that behavioural differences that correlate with morphological traits might promote adaptive mechanisms in simple environments, whereas neurologically based plastic behavior may be necessary to adapt in complex environments.

## Introduction

Species adaptation to the environment has been a revolutionary concept in the understanding of Biology since the publication of *On the Origin of Species* (Darwin, 1859). Gradual adaptations, often driven by the accumulation of mutations over many generations, allow physiological and morphological traits to evolve at the species level. However, evolutionary change can also occur rapidly, enabling organisms to adapt to sudden environmental shifts, such as those caused by climate change, or to establish themselves in new regions. A notable example of the latter is the introduction of non-native species, often a byproduct of globalization, which has become a significant ecological and economic challenge (*e.g.* Bertelsmeier, 2021; López et al., 2023). To become established in a new area, introduced species must cope with a range of diseases, resources, competitors, and conditions. In the case of the alien invasive species, these adaptations not only allow them to survive in the new place, but to expand their range and colonize new areas (*e.g.* Prentis et al., 2008; Stapley et al., 2015).

One of the mechanisms that enable species to rapidly modify the way they interact with the environment is through their behaviour (reviewed in Snell-Rood, 2013). At the population level, these behavioural adaptations can arise from two different mechanisms. One of them consists of the difference of interindividual behavioural patterns (personality; Dall et al., 2004) and their match with the environmental conditions, which determines the fitness of the individuals. Another mechanism consists in the intraindividual changes in behaviour through processes and experiences (plasticity; Dingemanse et al., 2010). While these concepts may directly apply to solitary species, the proximate causes of behavioural variability and the fitness outcome of its dynamics differs in eusocial organisms. In such species, evolutionary pressures and selections do not act at the individual but at the colony level (*e.g.*, mole rats, Jacobs & Jarvis, 1996; ants, Delgado & Solé, 2000). On one hand, interindividual behavioural variability in eusocial organisms often relies on task speciation (Jeanson & Weidenmüller, 2014), which is frequently related with individuals’ age (polyethism; reviewed in Beshers & Fewell, 2001), a form of intraindividual behavioural plasticity through development (also see Jeanson 2019). On the other hand, eusocial behaviour is strongly influenced by colony-level dynamics, as new behaviours often emerge as collective phenomena (reviewed in Lihoreau et al., 2012; Navas-Zuloaga et al., 2022).

However, in the last years a growing number of studies are demonstrating the importance of interindividual behavioural differences (*e.g.* Maák et al., 2020; Trigos-Peral et al., 2023) and intraindividual behavioural plasticity (reviewed in Lucon-Xiccato et al., 2024) in the context of coping with environmental challenges at the individual level in eusocial species. Whether due to phenotype, task specialization or perceived experiences, proximal mechanisms such as neuroanatomical development, are responsible for the variability of insect behavior. Although insect morphology stops changing after metamorphosis, their brains show large plasticity in the adult stage. For instance, the relative volumes of certain neuronal regions (*e.g.* Gronenberg et al., 1996) or even the total volume of ants’ brains can increase through maturation and experience in the adult stage (*Camponotus* sp.. Gronenberg et al., 1996; Yilmaz et al., 2016b; *Oecophylla smaragdina* and *Formica subsericea*, Kamhi et al., 2017). In polymorphic species, larger workers show larger optical lobes - necessary when foraging - and smaller workers show higher ratio of collars of the mushroom body to optic lobes, *i.e.* more processing than perception of the visual cues (Arganda et al., 2020). The navigation under natural skylight involves the increase of the synaptic buttons of visual projection neurons and volume of the central complex in *Cataglyphus nodus* (Grob et al., 2017). When learning, *Acromyrmex ambiguus* shows a temporal increase in the synaptic cluster (microglomureli) in the olfactory region rather than in the optic region of the mushroom body (Falibene et al., 2015).

The production, development, and maintenance of neuronal plasticity (*e.g.* new or larger neural routes) involves resource allocation and costs for the individual (Jeanson & Weidenmüller, 2014). Behavioural processes such as memory and learning allow faster adaptation (Fawcett et al., 2013) but involves the organisation of the brain functional areas (neuropils) in charge of processing sensory information (mushroom bodies of insects; Aso et al., 2014; Li & Strausfeld, 1999). Some of these costs could scale to the progenitor level, which might invest more resources to allow individuals coping with neuroanatomic plasticity, leading to a trade-off related with the quantity and quality of the offspring production (Clutton-Brock et al., 1985; Einum & Fleming, 2004.). Therefore, we expected an inverse relationship between the behavioural plasticity and the colony size on eusocial animals, *i.e.* queens investing in workers’ quality in small colonies but in workers’ quantity in large colonies (Kramer & Schaible, 2013). Under this scenario, large colonies might rely more on the polyethism and the behavioural traits arised by the collective than in the individual behavioural properties.

The Argentine ant (*Linepithema humile*, Mayr 1868) could be one of the most representative examples to illustrate trade-offs between the benefits and costs of behavioural variability due to the millions of members that form its supercolonies (non-competitive colonies; reviewed in Moffett, 2012) and their necessity to adapt to the introduced areas in which they behave as invasive. We therefore used the Argentine ant to highlight the individual capacities to interact with the environment and its dependence on neuroanatomy. At present, we do not know how differences and neuroethological plasticity of workers cause or affect colony adaptation in invasive species. We expect that in a polygynous superorganism (Hölldobler & Wilson, 1990) in which workers are almost non-genetically related (Keller & Fournier, 2002) and with the dimension of the Argentine ant introduced supercolonies (reviewed in Angulo et al., 2024), the behavioural variability should rely more on genetic or developmental differences among and within the supercolony members than in the individual capacity to modulate behavioural responses. On the contrary, some characteristics of the *L. humile* workers such as the rapid consolidation of long-term memory (Wagner et al., 2023) and traits found in other ant species like learning (Czaczkes et al., 2013; Jeanson & Weidenmüller, 2014), personality and tool use (Maák et al., 2020) or task syndromes (Trigos-Peral et al., 2023), suggest a high relevance of individual cognition in problem-solving in eusocial organised species (also shown in bees, Lemanski et al., 2021; bumblebees, Hill et al., 2023). Unraveling the underlying mechanisms involved in resolving environmental challenges faced by the introduced *L. humile* supercolony may help to prevent and control the continued expansion (reviewed in Angulo et al., 2024) and long-term stabilisation (Baratelli et al., 2023) of this ant species.

We performed standardised laboratory experiments with different environmental complexities to analyse the behavioural strategies used by the Argentine ants. We considered exploratory behaviour as a basic mechanism to solve problems in new environments (open field test, response to an object and solving maze). We focused on the relationship between the individual neuroanatomy and the behavioural plasticity in the face of environmental challenges to understand potential dependences of behaviour on the brain and thus, colony costs associated with individual production. Argentine ants have managed to spread widely in their introduced areas owing to their high activity, population density, and capacity to monopolise larger foraging territories (Carpintero & Reyes-López, 2008; Human & Gordon, 1996), so we expected experimental ants to show high level of exploratory behaviour. As a possible strategy of reducing developmental costs in maintaining a large population (Sanmartín-Villar et al., 2021), we expected ants showing low inter- and intraindividual behavioural variability when increasing the environment complexity and sparse relationship between the neuroanatomy (microglomureli organisation and brain region volumes). In other words, we expected that the colony interaction with the environment will be a product of individuals’ trial and errors and without cognitive processes associated (Lihoreau et al., 2012).

## Material and Methods

### Sampling collection and laboratory maintenance

*Linepithema humile* colonies were collected in NW Spain between the 23^rd^ and 28^th^ of August 2022 in five locations separated by ca. 7 km (in latitudinal order; Catoira: 42°40’18”N, 8°43’41”; Rianxo: 42°38’43”, 84°9’14”; Carril: 42°37’14”, 8°46’21”; Chazo: 42°36’27”, 8°51’25”; Vilanova: 42°34’00”, 8°49’42”). Queens, workers, and larvae were transferred with the soil of their nests to plastic containers with the inner part of the walls covered by polytetrafluoroethylene (Fluon®) to prevent ants from escaping. The contents of each collected nest were spread in plastic pot bases (97 x 40 x 7 cm) also covered by fluon®. We added plastic tubes filled with water and cotton to provide humidity to the ants and food *ad libitum* following the recipe based on Bhatkar & Whitcomb (1970) and improved according to Csata *et al*. (2020). This food was used throughout the experiment. Over the course of five days, all ants successively entered into the tubes and were transferred to new plastic containers and transported, without soil, to the Museum and Institute of Zoology of Warsaw (Poland). Ant nests (“source nest” hereafter) were maintained in temperature chambers at 26°C with a 12:12h light:dark period, the optimal temperature from egg to adult development of *L. humile* (Abril et al., 2010). Food was replaced in the source nest three times per week until the end of the experiment.

To control the age of the focal individuals and avoid behavioural and neural differences due to polyethism (*e.g.*, Fahrbach et al., 1998; Kamhi et al., 2019; Tripet & Nonacs, 2004) and different experiences (*e.g.*, Weidenmüller et al., 2009), we followed the development of last instar larvae until their metamorphosis. To facilitate this process and to standardise the experiences lived by focal individuals in their larval stage, we created homogeneous experimental groups by transferring from the source colonies 20 workers and five last-instar larvae to Petri dishes (Ø= 5.7 cm) containing a 1.5 ml Eppendorf tube filled with water and cotton and the tube’s lid filled with food. Twenty experimental group replicates were initially isolated from each source nest. In cases where two callow workers emerged in the same group, the newest callow worker was transferred to fresh experimental groups. We reached a maximum of 50 and minimum of 20 experimental groups for the source nests. We checked each experimental group daily for new emergences. One day after emerging, focal individuals were marked with pen markers (Edding Creative 751©). Focal individuals were easily identified even one day after their emergence because the colour of their cuticle was lighter than the one of their nestmates. Behavioural tests started ten days after the emergence of focal individuals, a period that we considered optimal to guarantee the development of the locomotive workers’ function (see also Abril et al., 2010).

### Behavioural test

To study inter- and intra individuals’ behavioural variability, we exposed each focal individual (N= 172) to a series of three consecutive 10-minute tests. Five observations of the three different tests, one per day for five consecutive days, were made (N= 860 tests) at 23.75±0.73 °C and 54.67±2.78 % of humidity, testing each individual over a similar time frame. Tests were always conducted in the following order for each individual and observation: (i) open field test, focused on the percentage of new area explored; (ii) object exposure test, focused on the interaction with new objects; and (iii) maze test, focused on the decision making on whether to explore further in a complex environment. A total of 13 individuals were not exposed to the objects and mazes, *i.e.* they remained for 30 min in the open field (see below), to control the effect of the exposure to new objects on the behaviour and brain structure. A total of 20 individuals were not exposed to any of the behavioural tests as a control for possible inherent factors of our procedure such as manipulation and isolation (Bernadou et al., 2018; Seid & Junge, 2016) that could cause some changes to the brain.

Behavioural tests were performed in separate Petri dishes inside three 60 x 70 x 60 cm wooden boxes with white walls to limit visual references and thus, without distal clues for the focal ants (see Harris et al., 2010). Ants’ behaviour was recorded from the top of the boxes with cameras (two Sony FDR-AX53; one Canon Legria HF M56) and tracked with AnimalTA software (Chiara & Kim, 2023). A maximum of 10 ants were recorded at the same time for a single camera. Before each series of tests started, we transferred the experimental groups from the temperature chambers and placed them in the wooden boxes for 10 minutes as pre-acclimatisation.

#### Open field test

Following the 10 min of acclimatisation, focal ants were carefully isolated into their respective empty Petri dishes (Ø= 5.7 cm) using a brush. Petri dishes were covered with their lids after the addition of each individual. These dishes possessed a red circle of 1 cm diameter drawn on the centre and outer part of the bottom dish as a reference for placing the object in the following test (see next paragraph). We assumed that ants did not perceive this circle due to its position (no odour because it was out of the dish). Recordings started before the introduction of the first ant and lasted 10 min after the introduction of the last ant in the experimental setup. By doing so, we were able to register the behaviour of all focal individuals since their contact with the new arena. This procedure was also applied to the other behavioural tests. Upon introducing the last ant, we closed the wooden boxes with a lateral white wooden cover.

#### Object exposure test

Following the 10-minute open field test, we removed the cover of the wooden boxes and introduced an object in each Petri dish. The objects were added in the centre of the arena, matching its contour with the red circle designed. After adding the object to the last dish, the cover of the wooden box was put back and we recorded the ants’ behaviour for 10 mins. Objects consisted of a three-level circular pyramid (Ø= 1 cm; Fig. 1A) designed on Tinkercad (https://www.tinkercad.com/) and printed on resin (Anycubic Eco UV Resin White) using Chitubox® and a 3-D printer (Elegoo Mars 2P). The objects were painted in blue (Pinty Plus Aqua; Ref. 320). We consider that the shape, material, colour, and odour were uncommon in the focal species environment and thus, relevant as a novelty stimulus. After the end of each test, the objects were washed with hand soap and tap water and allowed to dry under sunlight.

**Figure 1.**
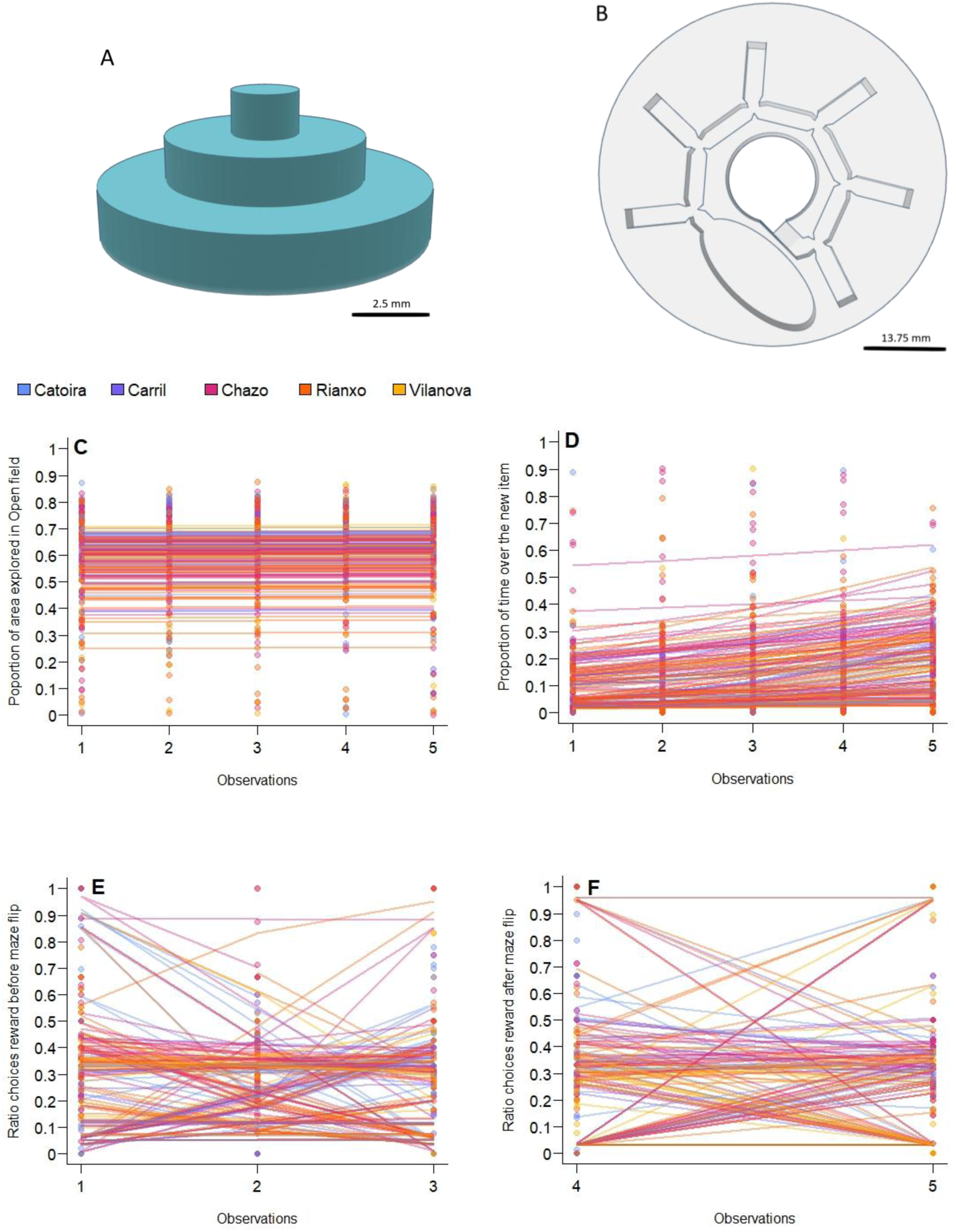
Behavioural tests and patterns of *L. humile* workers (regression models) across observations. **A-B**: Schematic representation of the novel object (A) and maze (B) used in the behavioural tests. Both objects can be downloaded for printing from https://www.tinkercad.com/things/2lNxDsoOoAD-neophobia-object-miiz and https://www.tinkercad.com/things/7IELVyTIwXr-maze-miiz. **C**: Exploration in the open field test. **D**: Interaction with the object. **E**: Relative number of times ants selected the path connecting to the reward instead of the dead-end paths of the maze (*Ratio choices reward*) before maze direction flip. **F**: Relative number of times ants selected the path connecting to the reward instead of the dead-end paths of the maze (*Ratio choices reward*) after maze direction flip.

#### Maze test

Following the 10 min object exposure test, ants were individually transferred to a new Petri dish (Ø= 5.7 cm) respectively containing a white resin-printed maze (height 4 mm, Ø= 5.7 cm; Fig. 1B). The maze consisted of a circular platform with two inner chambers (start and end) connected by a concentric path of 3 mm wide. The path was composed of a sequence of five 1 cm segments. Each segment ended in a Y-choice with the next two branches oriented 130° from the previous path. Opposed at an angle of 100° to each of the segments, the maze possessed a dead-end 1 cm segment. In this way, the maze was composed of six Y-choices connected by the concentrical path. Half of the mazes had dead-ends directed toward the left and the other half had dead-ends directed toward the right. On each Y-choice, ants had the choice between selecting a dead-end path or to continue advancing through the maze. The three vertices of the Y-choice were expanded to the interior of the path as projections constraining it to 2 mm. This forced the focal ants to face the two possible choices and avoid them selecting a choice based on the possible thigmotaxis (wall-following) used to guide them through the previous path (see Czaczkes, 2018). Focal ants were introduced to the ovoid start chamber of the maze (2.1 x 1 cm) with only one exit towards the maze path. We covered the maze with a plexiglas layer to avoid ants escaping. Ants that selected six times the same direction following the concentric path through the six consecutive Y-choices encountered the ending reward chamber. This reward chamber was circular (Ø= 1 cm) and positioned in the middle of the Petri dish (occupying the same place as the red circle of the open field test and the object on the object exposure test). Before each test, a drop of sucrose was added in this reward chamber (1M; see Czaczkes, 2018). The ending chamber was the only section of the maze without resin floor, to allow a better removal of the sucrose drop. The sucrose was added to stimulate the ants that found the end chamber for future trials, although our aim was not to test learning (no training).

To analyse the variability of choice strategy under novel environments, focal ants were exposed on the three first consecutive observations to mazes with a concentric path oriented in one direction and to mazes oriented in the opposite direction on the last two observations. By this, we could check if ants were habituated to the maze and how they reacted after a change, which we assumed is a more demanding cognitive process (central integration; see Menzel & Giurfa, 2001) and thus, the opportunity to compare personality traits and cognition based on different mechanisms (see Carere & Locurto, 2011). After 10 min of recording, ants were returned to their original experimental group and then to the temperature chamber until the next day, in which tests were repeated.

Ants belonging to the control group were maintained 30 minutes in empty arenas (open field) to control the effect of experiencing new or/and complex environments (object, maze) while considering the effects of experimentation (manipulation and isolation). Similar to the experimental group, we made 5 observations over 5 consecutive days.

After 16 days of their emergence (immediately after the last test on the 5^th^ day for focal ants), individuals were isolated in 15 ml falcon tubes and placed in a thermally isolated box with ice packs until brain extraction.

### Brain extraction and fixation

Brain extraction was done following the protocol of Ott (2008). Focal and control ants (N=205) were anaesthetised on ice. Immediately after decapitation, the eye span of each ant was measured using a micrometre under a binocular lens (Olympus SZH10). Brain extractions were performed in HEPES solution (150 mM NaCl, 5 mM KCl, 5 mM CaCl₂, 25 mM sucrose, 10 mM HEPES) in silicone-coated Petri dishes. Extracted brains were individually fixed in 1% Zn-PFA solution overnight and continuously shaken (60 rpm; Thermo Fisher) at room temperature. The brains were then rinsed in HEPES solution six times for 10 minutes each. Subsequently, the brains were fixed in Dent’s Fixative (1:4 DMSO-Methanol solution) at room temperature for 1 hour and 30 minutes. After fixation, the brains were stored in 100% methanol at -20°C until the immunostaining procedure. Once all brains were collected, they were shipped to the University Rey Juan Carlos (Spain) under cold conditions to ensure preservation.

### Immunohistochemical analysis and imaging

To image the neuroanatomy of the extracted brains, they were rehydrated in 0.1 M Tris buffer and seated in a block and permeabilization solution of normal goat serum (NGS) for 1 hour (PBSTN solution: 5% NGS + 0.005% sodium azide in 0.2% Triton-X phosphate buffered saline [PBST]) at room temperature on a shaker. After this step, brains were incubated on the shaker during four nights at room temperature in the primary antibody solution (SYNORF1 – AB_2315426, Developmental Studies Hybridoma Bank – diluted 1:30 in PBSTN). Then, brains were washed six times (10 minutes/wash) in 0.2% PBST and incubated for another three nights protected from light on a shaker in AlexaFluor488 (ThermoFisher) Goat anti-Mouse IgG (H+L) Cross-Adsorbed ReadyProbes™ Secondary Antibody (1:500 in PBSTN). Once immunolabeled, brains were washed again for six times in 0.2% PBST (10 minutes/wash) and dehydrated in a series of ethanol:PBS solutions (10 minutes each in 30%, 50%, 70%, 95%, 100%, 100% solutions). Then, they were either stored overnight at -20°C or directly mounted as follows. To mount the brains, they were cleared in methyl salicylate (removing the ethanol solution) and seated in wells of stainless-steel slides containing methyl salicylate. Brains were imaged on an Olympus FV-3000 laser scanning confocal microscope. Whole brain images were taken with either a 20X (NA = 0.8) or a 40X (NA = 0.95) objective, in z-steps of 2 μm thickness. Close ups of the right mushroom body calyces (both lateral and medial) were taken with an oil immersion 100x objective, in z-steps of 0.5 μm thickness. We selected regions where the mushroom body calyces and the peduncle was visible.

### Neuroanatomic analyses

#### Volumetry

We aimed to quantify relative volumes of the following functionally distinct brain neuropils (Ito et al., 2014): the optic lobe (with lamina excluded - because it was not always present after dissection), the antennal lobe, the central complex (including central body, protocerebral bridge and nodes), and the mushroom body, where both the medial and lateral calyces, the peduncle, and lobules were separately identified. To avoid possible human bias and to save labelling time, a statistical template of *L. humile* brains was created by co-registering ten of the extracted brains. For these ten brains, the mentioned functional neuropils and the rest of the undifferentiated central brain were manually labelled in Amira (FEI – Amira 3D 2022.2) in the right hemisphere of each brain (only the central complex was labelled in both hemispheres, but only half of its volume was considered in the analyses). These manually obtained labels were used to create consensus labels in the template, allowing for automatic labelling of the rest of the registered brains. Automatic labels were smothered with Amira and manually corrected when needed. To calculate relative volumes of each neuropil, we calculated the volume of each region and of the whole brain in Amira (module Material Statistics) and computed the ratio between them. The ratio between primary (sensory) and secondary (processing) neuropils was calculated by dividing the volume of the optic lobes and antennal lobes by the volume of the mushroom bodies and the central complex.

#### Template construction and automatic labelling of brains

Following Arganda-Carreras et al. protocol (2017, 2018), the *L. humile* brain template was created using diffeomorphic space. Diffeomorphisms, which are smooth and invertible mappings of 3-D images, are commonly utilised in computational neuroimaging due to their topological properties that facilitate brain co-registration and the measurement of structural variation (Arganda-Carreras et al., 2017; Klein et al., 2009). The brain template was constructed from 3-D average-shape brains derived from the co-registration of 10 original brains obtained through confocal scans.

Initially, variations in illumination among the original brain images were corrected using histogram matching (Yoo, 2004). Co-registration involved an initial affine transformation to maximise mutual information between brain volumes, identifying common neuropil shapes. This was followed by iterative refinement using local non-rigid transformations to maximise the cross-correlation of voxel intensities among the co-registered brains. The final average shapes were generated by computing the voxel-wise median of the co-registered brains.

After creating the anatomical template, we applied the same diffeomorphic transformations to each label image of the original 10 brains, as performed on their corresponding brain anatomy. This was followed by per-voxel majority voting across all deformed label images of the same brain to produce consensus labels.

To automatically label new brain images, we registered individual brain images against the template using the same two-step method described above: an initial affine registration maximising mutual information, followed by a non-rigid registration optimising cross-correlation.

The inverse transformations were then applied to the template’s regional labels, automatically assigning label values to the individual grey value brain images registered against the template.

All steps were implemented using the Advanced Normalisation Tools (ANTs) software (Avants et al., 2011) after converting grey and label images to the open NRRD format in Fiji (Schindelin et al., 2012).

#### Microglomeruli

Microglomeruli numbers were counted manually in close-ups of the right mushroom body calyces using Amira. Close-up images of the right mushroom body calyces were analyzed to manually measure the number of microglomeruli. Using Amira, three cylindrical regions (each spanning 5 μm along the z-axis) were selected for analysis within each calyx: one in the collar and two in the lip. For the lip, regions were differentiated into the non-dense (volume: 256.288 μm³) and dense areas (volume: 97 μm³). The medial region of the calyx was located using the pedunculus as a reference, and circles were drawn on ten consecutive slices in this region.

Microglomeruli contours were analyzed in slice 5 of the ten marked slices. A masking threshold of 700 was applied in Amira to identify microglomeruli contours, and the generated labels were smoothed using the Smooth Label tool (size: 3). This step helped standardize the contours of the microglomeruli within the cylindrical regions.

Images of slice 5, with the contours generated in Amira, were then imported into ImageJ for counting. Using the Multi-Point Tool in ImageJ (Schindelin et al., 2012), microglomeruli were marked within the defined contours. For validation, 3-D images in Amira were reviewed to confirm that all labeled contours corresponded to microglomeruli.

### Statistical analyses

All analyses were performed using R version 4.4.2 (www.r-project.org). Assumptions of normality were checked for every variable using the *descdist* function. Non-parametric analyses were used instead of data transformation. When comparing means, we used ANOVA (*aov*), Kruskal-Wallis (*kruskal.test*), or Wilcoxon tests (*wilcox.test*) considering the dependency of the variables. Mean values are shown with standard errors (x_±SE). When comparing regression lines, we used Linear Mixed Models (LMMs) for models following Normal distribution (*lmer*) and Generalised Linear Mixed Models (GLMMs) using Template Model Builder (*glmmTMB*) for those following Negative Binomial (left-skewed and overdispersed continuous events) and Beta distribution (skewed continuous proportions variables ranged between 0 and 1). Knowing the dependency on body size of locomotion in ants (*e.g.*, Hurlbert et al., 2008), we used eye span as a proxy of body size as a covariable for all the analyses. We tested the dependency of the measured behavioural variables on body size (covariable) and across observations (fixed factor). We added the nest source and the ant worker identification code as random factors to take into account the potential variability among workers from different colonies (*e.g.* see colony personalities in Pinter-Wollman, 2012; behavioural differences in the populations surrounding the selected populations in Sanmartín-Villar et al., 2022) and to avoid pseudoreplication (individual repeated measures across observations).

To analyse the underlying mechanisms driving the measured variables on the three behavioural tests, we first analysed the effect of the cited factors on the locomotive traits: the average moving speed (calculated by discounting stationary events) and the proportion of time moving on each observation. All the tracks were smoothed on AnimalTA with Salvitzky-Golay filter with window length 15 and polyorder 2 to reduce the noise of the tracks. We considered the ant moving if its speed was above 2mm/s.

We used exploration (percentage of the total area visited in 10 min) as a variable to analyse the ant efficiency (or willingness) (i) to cover an unknown area (open field test), (ii) to cover a known area (after 10 min of acclimatisation) under the presence of a new object (object exposure test), and (iii) to discover unvisited paths (maze test). Based on the measurements of 25 ants’ sizes, we considered that ants were exploring an area of 5 mm^2^ around them for these exploration analyses. The exploration and the use of the inner part of the arena (38.48 mm^2^) were compared between experimental and control individuals by comparing the variables obtained in the open field (10 first min of observation) and the exposure to the object (10 next min) with the first 10 min and next 10 min of observation in a control open field.

We used the proportion of time spent interacting with the object as a proxy of neophobia/neophilia. We used the relative number of choices allowing maze exploration (those following the concentric path; after now, “ratio of choices to reward”) as a proxy of ant exploration in complex scenarios. We consider that the ant made a choice if it crossed both the projections on each Y junction of the maze. To calculate the ratio of choices to reward, we first divided the total number of choices allowing maze exploration by two to avoid considering the data of individuals crossing backwards a choice point. In the case of obtaining decimal values, the numbers were rounded to the next superior complete digit. For instance, a value of three units meant that one ant crossed three times the same choice, two forward and one backwards. By dividing by two (3/2= 1.5) and rounding to the next superior complete digit we obtain the number of forward crosses (2). Later, we divided the outcome variable by the total number of choices (total number of choices allowing maze exploration plus the total number of choices driving to a dead-end segment). We analysed the direction (left or right) of the first path selected in the maze to disentangle if this selection was conditioned by (i) lateralization preference (*e.g*. trend to select left or right paths; see Wystrach et al., 2016); (ii) previous choices performed in the previous observation (*e.g.* starting to the same direction as in the previous observation); (iii) the openness of the path (assuming that focal ants were able to perceive the end of the path in the selection moment, they could have a preference to select a path that allows continuing exploring or going to a dead-end path).

To analyse if individuals showed different behavioural patterns across time (intraindividual differences in behavioural plasticity) and if these patterns differed among individuals (interindividual differences in behavioural plasticity) we compared the AIC values of models with and without observations as random slopes for each individual intercept (see Sanmartín-Villar et al., 2021). AIC values are shown as the difference (Δ) between AIC_without random slopes_ - AIC_with random slopes_. Models with random slopes were only selected under ΔAIC positive values higher than 2 units (the rule of thumb when selecting AIC models, *e.g.* Mazerolle, 2006). The marginal (R^2^m; variance explained by the fixed factors) and conditional (R^2^c; variance explained by the full model) coefficient of determination are given for the most explanatory model (lower AIC value).

To analyse the consistency of behaviours, we analyse unpredictability, repeatability, and coefficients of variation. We calculated unpredictability following the residual individual standard deviation (riSD), a standard deviation that “estimates variation around the expected values at every time step instead of estimating the amount of variation around the average of all of the observations” (Stamps et al., 2012). We selected this method among others (see Cleasby et al., 2015) because its simplicity and the similarity of the produced results respect other methods that we found in previous experiments (e.g., Sanmartín-Villar et al., 2021).

We calculated repeatability using the *rptGaussian* function (*rptR,* Stoffel et al., 2017; Suppl. Material), which is based on the Intraclass Correlation Coefficient (see Nakagawa & Schielzeth, 2010). Due to criticisms of using this coefficient without taking into account the measurement errors (see Dingemanse et al., 2022), we also calculated the coefficient of inter- and intraindividual variation (formulae 4 and 5 in Dingemanse et al., 2022) for the behaviours that we consider more explanatory of each test (exploration in opened field test, object interaction in object exposure test, and the ratio of choices to reward in the maze test).

Behavioural syndromes (the correlation of personality traits; Bell, 2006) were analysed by adding one behavioural measure as variable and other as covariable of GLMMs. We compared the correlation between exploration in open fields and the interaction with the new objects and both of them with the maze participation (ant willingness to enter into the maze after being added in the starting area), the ratio of choices to reward, and the number of times that ants reached the end of the maze.

To test the correlation between behaviours dependent of different information processing (see Carere & Locurto, 2011), we compared the proportion of time spent on the object for the first time (neophilia/neophobia) with the total number of choices and the ratio of choices to reward in the fourth maze test (maze performance after maze flip).

The behavioural relationship with neuropils relative volume and number of microglomeruli was analysed using MANOVA tests (*manova*), adding the behavioural means or variances as dependent variables and the neuropils relative volume or the number of microglomeruli from the measured brain areas as independent variables. We analysed by the same method the effect of worker size on the neuropil’s relative volumetry and number of microglomeruli. We also performed Principal Components Analysis (PCA) to analyse which variables explain most of the variance in behaviour, neuropils relative volume and number of microglomeruli. We then used the resulting PCs to compare behaviour with the two brain traits. We could not compare the PCs from the two brain measures in the same analyses because of the quality of the images (some good for volume analyses but not for microglomeruli and *vice versa*).

## Results

### Behavioural variability

#### Test performance and individual differences

Body size affected locomotive traits. Larger ants were faster and spent more time moving in the behavioural tests (Table 1). The locomotive traits changed across observations. The ants’ average speed decreased across observations, but the proportion of time spent moving showed different patterns across time: they did not change in the open field, increased in the object exposure test, and decreased in the maze (Table 1).

**Table 1.**
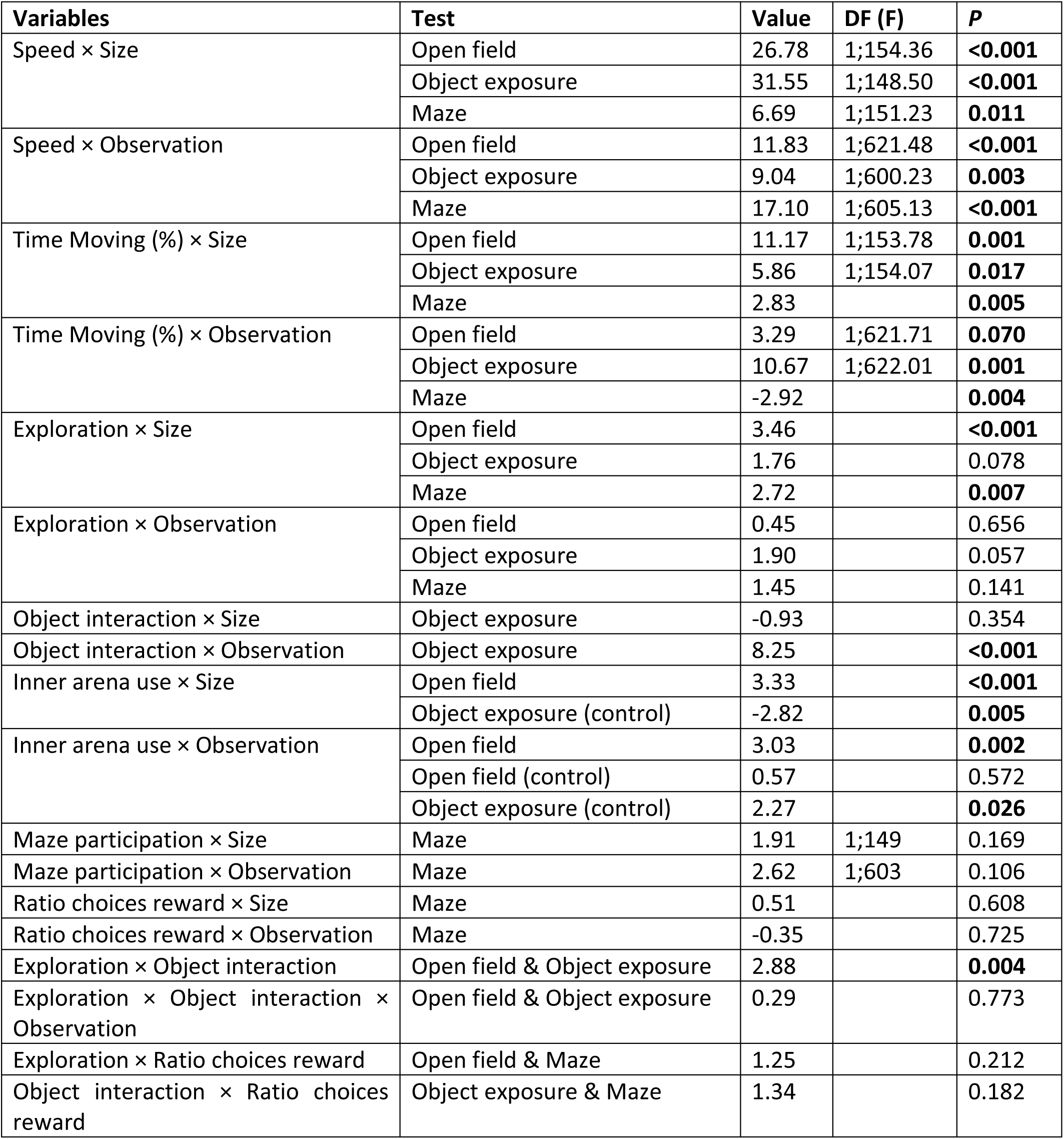
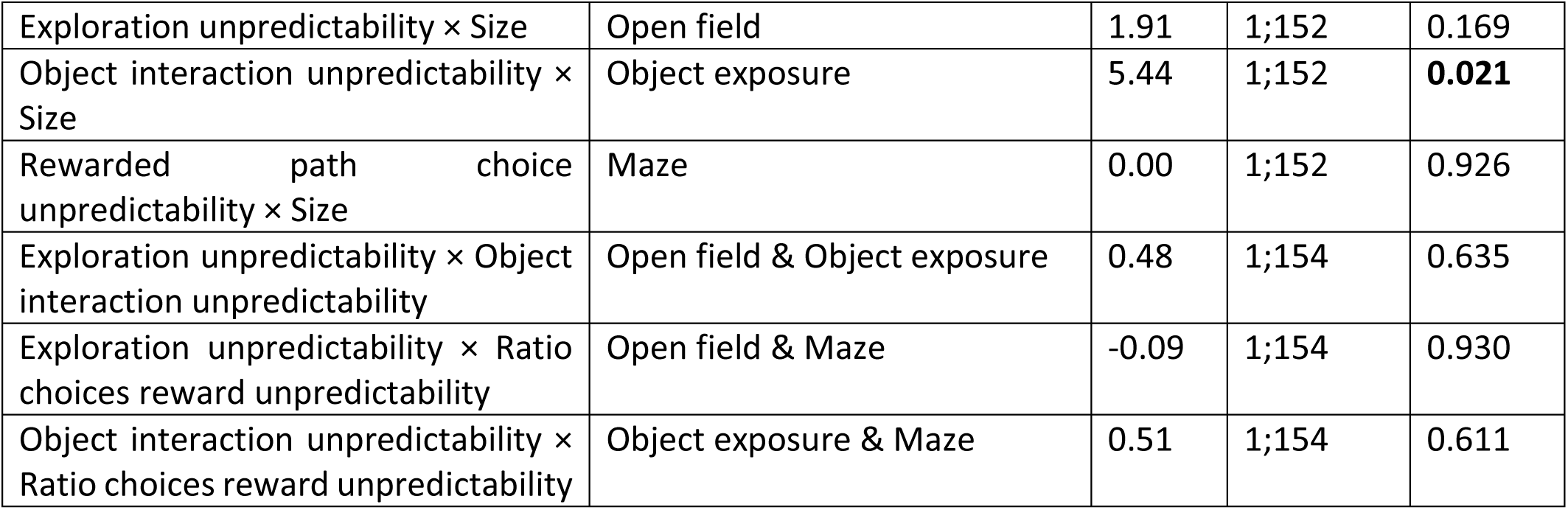
Statistical values (*Value*: t, F or Z) for the behavioural variables measured. *Ratio choices reward*: relative number of times ants selected the path connecting to the reward instead of the dead-end paths of the maze.

Workers’ exploration depended on body size but showed similar and consistent interindividual patterns across time. Larger ants explored a larger proportion of area in the open field and the maze, but arena exploration was not dependent on ant size in the object exposure test (Table 1). Individual’s exploration pattern was constant through observations (no intraindividual differences) in the open field (60.60±0.69% of the area) and maze (26.81±0.74%) but it showed a marginal increase in object exposure tests (1^st^ observation: 53.22±1.71%, 5^th^ observation: 57.71±1.67%; Table 1). In the three behavioural tests, the exploration pattern through observations showed no differences among individuals (no interindividual differences; Table 2; Figure 1C).

**Table 2.** Intraindividual differences. *Ratio choices reward*: relative number of times ants selected the path connecting to the reward instead of the dead-end paths of the maze.

| Variable | Test | $\Delta AIC$ | $R^2m$ | $R^2c$ |
| --- | --- | --- | --- | --- |
| Exploration | Open field |  | 0.06 | 0.49 |
|  | Object exposure |  | 0.01 | 0.11 |
|  | Maze |  | -0.22 | -2.82 |
| Object interaction | Object exposure | 9.02 | 0.16 | 1.60 |
| Ratio choices reward before flip | Maze | 73.55 |  |  |
| Ratio choices reward after flip | Maze | 52.72 |  |  |

The addition of the object represented a change in the exploration of the arena’s areas and its interaction highlighted different behavioural patterns. Smaller ants used more the inner zone of the experimental dish when the object was absent (open field and control group in the object exposure test) but body size did not affect the interaction with the object in the object exposure test (Table 1). On the first observation, ants were more attracted to the new object (9.84±1.21% of the time interacting) than to the inner area of the dish in the open field during the 10 previous minutes (5.67±0.97%; V= 7816, *p*= 0.003). The proportion of time spent in the inner zone of the dish by the control group (object absent) was even lower (2.73±1.58%), but no statistical differences were found with the experimental group due to the sample size (W= 686, *p*= 0.258). The time spent interacting with the object increased through observations (5^th^ observation: 18.28±1.24%; F= 25.13, *p*< 0.001). This increase was more pronounced than the increase of the use of the inner zone in the open field or for control ants in the object exposure test (object absent; 5^th^ observation < 6.17±10.82%). The increase of the proportion of time interacting with the object across observations differed among ants (interindividual differences; Table 2; Figure 1D). Unexpectedly, ants exposed to the object (focals) modified the use of the inner zone of the dish in the open field (object absent) through observations, while those never exposed (controls) did not change it (Table 1). After 10 min of test, *i.e.*, in the object exposure test, control individuals (object absent) also increased the use of the inner zone of the dish though observations (Table 1), although this increase was lower than the observed in the experimental group and in the open field test.

Focal workers showed different strategies toward the rewarded path. The ratio of choices for rewarded path differed among focal workers (interindividual differences) both before (28.26±1.16%; Figure 1E) and after flipping the maze direction (27.51±1.56%; Figure 1F). Individual choices followed such different trends that it was not possible to statistically detect an overall effect across observations (F< 0.20, *p*> 0.689). Ant size did not affect the ratio of choices for rewarded path (Z= -0.35, *p*= 0.725). More detailed information about maze performance and strategies of path selection are explained in Supplementary Material.

#### Behavioural syndromes

Behaviours measured in different tests were correlated within individuals. Ants that explored more in the open field spent more time interacting with the new object (Table 1). This correlation was not affected by the observation number (Table 1). Ants that explored more in the open field participated more in the maze exploration (t= -4.37, df= 149, *p*< 0.001) and reached more times the reward (ꭓ^2^_2_= 18.18, *p*< 0.001), but they did not reach faster the reward of the maze after having reached it once (Z= 0.61, *p*= 1). Ants’ exploration or object interaction were not correlated with the ratio of choices to reward (Table 1).

We found no behavioural syndromes involving different information processing. The first interaction with the object was not correlated with maze participation (F_1;149_= 0.075, p= 0.785) or with the ratio of choices to reward of the fourth observation (first encounter of the flipped maze) (S= 627247, p= 0.915, r= 0.01).

#### Unpredictability

Focal workers behavioural unpredictability was independent in the three behavioural tests, and it was not affected by their behavioural performance. No correlations were found between the unpredictability of the exploration in open field, object interaction, and the ratio of choices to reward (t< [0.51], df= 154, *p*> 0.635). No differences in unpredictability were found between control and focal individuals for their exploration in the open field (F_1;165_= 0.12, *p*= 0.731). The introduction of a new object decreased the unpredictability of the ants’ use of the inner part of the arena when the object was present - or after 10 min of test for the control group - (control-focal: F_4;165_= 4.75, *p*= 0.031), while the unpredictability to use the inner part did not change in the exploration on the open field test (F_4;165_= 0.12, p= 0.731). No differences in unpredictability for the object interaction were found for individuals increasing or decreasing the interaction over the observations (t= -0.53, df= 154, *p*= 0.592). The unpredictability of the ratio of choices to reward was similar for those individuals increasing or decreasing this ratio considering all observations or only those before maze flip (t< 1.36, df= 137.04, *p*> 0.177). Ant’s exploration and the ratio of choices to reward were similarly unpredictable according to the ant size (Table 1), but the interaction with the new object was less unpredictable in larger ants (Table 1).

#### Coefficients of variation

The behavioural variability observed among focal and control ant workers was observed in the object exposure tests, but not in the open field. The coefficient of intraindividual variation for exploration was similar among control (21.52±3.23%) and focal ants (23.80±1.09%; F_1;165_= 0.30, *p*= 0.586). No differences were found for worker size (t= -1.63, *p*= 0.193). The coefficient of intraindividual variation for the interaction with the inner area differed between control (without object; 0.91±0.09%) and focal ants (with object; 13.67±0.80%; F_1;164_= 5.40, *p*= 0.021) in the object exposure test. Smaller ants showed higher variation (t= 12.70, *p*< 0.007). The coefficient of intraindividual variation for the ratio of choices to reward (23.06±0.92%) did not depend on worker size (t= -1.41, df= 154, *p*= 0.162).

The coefficient of interindividual variation was similar among the control (33.35%) and the focal group (20.03%; Z= 1.31; critical value= 1.96) for exploration but strongly differed for the interaction with the object (control: 33.16%; focal: 75.57%; Z= -3.63; critical value= 1.96). Ants showed an interindividual variation of 43.17% for the ratio of choices to reward.

### Neuroanatomy

#### Volumetry

The relative volume of the cited functional neuropils was only analysed in 38 focal workers (daily exposed to the three behavioural tests over five days) due to logistics. Unfortunately, the relative neuropils volumes of the control group (not exposed to behavioural tests) could only be analysed in two individuals and were therefore discarded.

The relative volumes of the different neuropils were uncorrelated in most cases. No significant correlation was found between the body size and the relative volume of the brain neuropils measured (F_1;30_ < 2.21, *p*> 0.148). The relative volumes of the medial and lateral mushroom body calyces were positively correlated (t= 5.09, df= 36, *p*> 0.001, r= 64.68%). The relative volume of the central complex and the optic lobe was negatively correlated (t= -2.39, df= 36, *p*= 0.022). No correlations were found between the relative volume of (i) the optic and antennal lobes, (ii) the central complex and the mushroom body, (iii) and the mushroom body and the sensory (optic or antennal) lobes (t< [1.37], *p*> 0.181, r< 22.19%).

#### Microglomeruli

The number of microglomeruli of the analysed regions (collar, dense lip, non-dense lip) were positively correlated (t > 3.75, *p* < 0.001, r > 54.65). The number of microglomeruli present in the collar showed a positive correlation with the body size (F_1;24_= 5.42, *p*= 0.029). No significant effect of worker size was found for the microglomeruli of the lip (F_1;24_< 0.69, *p*= 0.414).

### Behaviour-brain correlation

The relative volume of the neuropils conditioned the behavioural performance in complex environments. Ant workers with higher relative optic lobe volumes moved faster in the presence of the new object (F_1;29_= 7.43, *p*= 0.011; Figure 2A). Ant workers with lower primary:secondary neuropils ratio showed higher exploration when exposed to the object (F_1;29_= 4.84, *p*= 0.036; Figure 2B) and to the maze (F_1;29_= 5.30, *p*= 0.029; Figure 2C). No other significant correlations were found between the neuropils’ relative volume and the mean and variance of speed or exploration in the three behavioural tests performed (F_1;29_< 3.02, *p*= 0.093). The regression slopes of the ratio of choices to reward before the maze flip were negatively correlated with the relative volume of the medial mushroom body calyx (F_1;29_= 7.65, *p*= 0.010; Figure 2D; but not with the lateral calyx: F_1;29_= 2.68, *p*= 0.113) and positively correlated with the relative volume of the central complex (F_1;29_= 10.98, *p*= 0.002; Figure 2E). No significant differences were found when comparing behavioural and brain volumetry PCs (F_1;35_< 1.73, *p*> 0.196).

**Figure 2.**
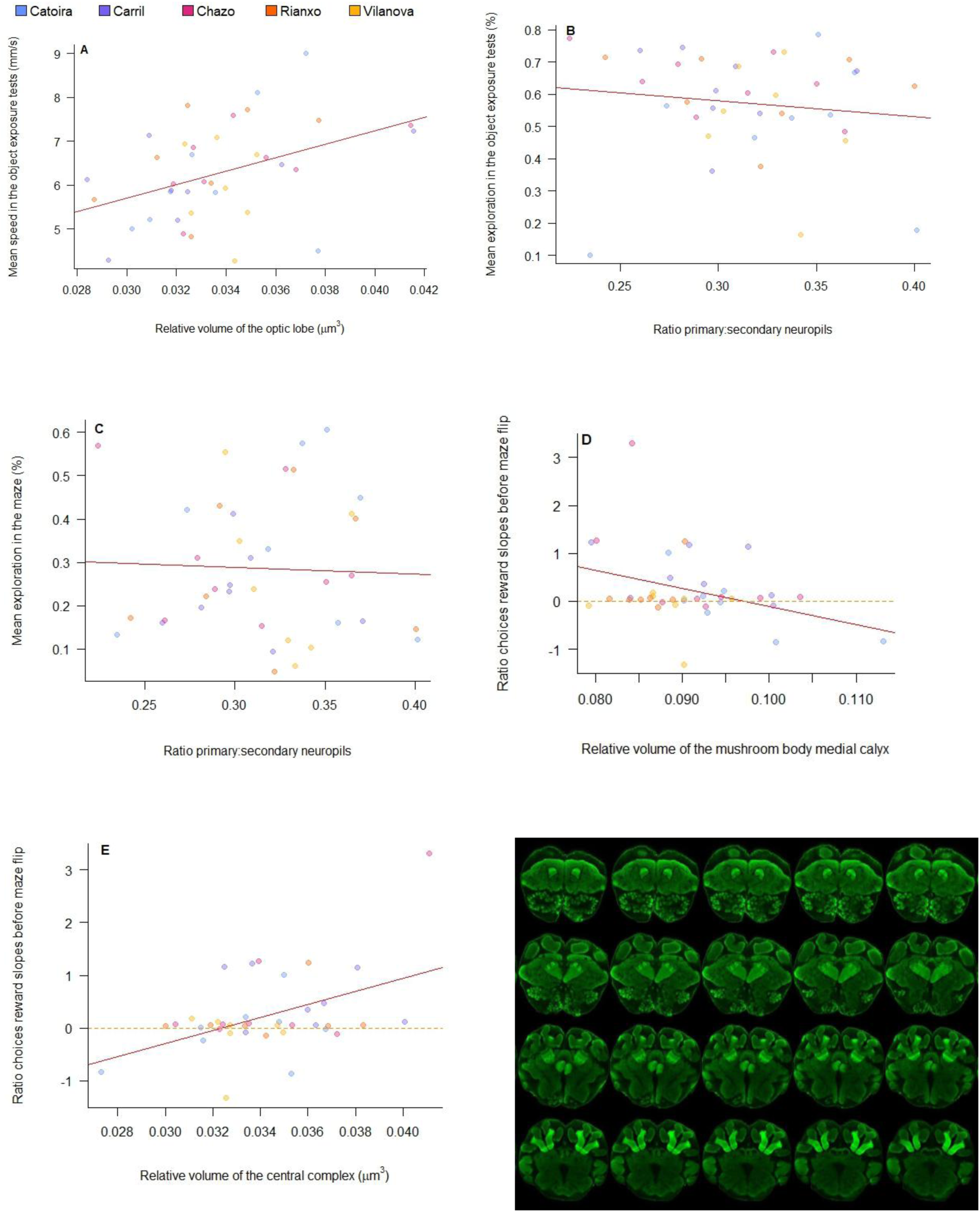
Correlation between brain volumetry and behaviour of *L. humile* workers: correlation between the relative volume of the optic lobe and the mean speed performed (**A**); correlation between the ratio primary:secondary neuropils and the mean exploration performed in presence of the object (**B**), on the maze (**C**), and the slopes for the relative number of times ants selected the path connecting to the reward instead of the dead-end paths of the maze (*Ratio choices reward*) before flip (**D**); correlation between the relative volume of the central complex and the slopes for the relative number of times ants selected the path connecting to the reward instead of the dead-end paths of the maze before flip (**E**). Continuous red: linear regression between the two variables. Dashed orange: zero value to facilitate the visual comparison of negative and positive slopes. F: different sections of a *L. humile* brain showing the neuropils analysed (synapsin in green).

The number of microglomeruli was similar among control and focal ants and among those showing different mean behaviours. No differences were found between the number of microglomeruli of individuals exposed (N= 38) or not exposed (control; N= 5) to behavioural tests in any of the three analysed brain regions (F_1;38_< 1.64, *p*> 0.209), but a trend suggested that control individuals showed higher variance than the focal ones in the number of microglomeruli present in the collar (F_1;38_= 3.90, *p*= 0.056). The number of microglomeruli of the three regions analysed was not correlated with the average speed (F_1;24_< 1.20, *p*> 0.334) or the average exploration (F_1;24_< 1.17, *p*> 0.345) performed on the three behavioural tests. Nor were correlations found with the mean object interactions and the mean ratio of choices to reward before and after maze flip (F_1;24_< 1.79, *p*> 0.189).

However, the number of microglomeruli differed among individuals showing different behavioural patterns under complex environments. The number of microglomeruli of the three regions analysed was not correlated with the variance of speed (F_1;24_< 1.20, *p*> 0.334) or exploration (F_1;24_< 1.17, *p*> 0.345) performed on the three behavioural tests. Individuals with higher number of microglomeruli on the dense lip showed higher variance (F_1;24_= 6.50, *p*= 0.018; Figure 3A), more unpredictability (riSD: F_1;24_= 7.87, *p*= 0.010; Figure 3B) on the object interaction, and higher increase of the ratio of choices to reward before maze flip (F_1;24_= 9.10, *p*= 0.006; Figure 3C). No correlations were found between the object interaction variance and the number of microglomeruli from the collar or non-dense lip (F_1;24_< 1.23, *p*> 0.278), nor between the slopes of the ratio of choices to reward before maze flip and the non-dense lip (F_1;24_= 1.01, *p*= 0.326). Individuals with higher numbers of microglomeruli in the collar showed decreased slopes of the ratio of choices to reward before maze flip (F_1;24_= 5.81, *p*= 0.024; Figure 3D). The variance after maze flip was not correlated with the number of microglomeruli in any region (F_1;24_< 0.85, *p*> 0.367). No significant differences were found when comparing behavioural and microglomeruli PCs (F_1;13_< 1.18, *p*> 0.339).

**Figure 3.**
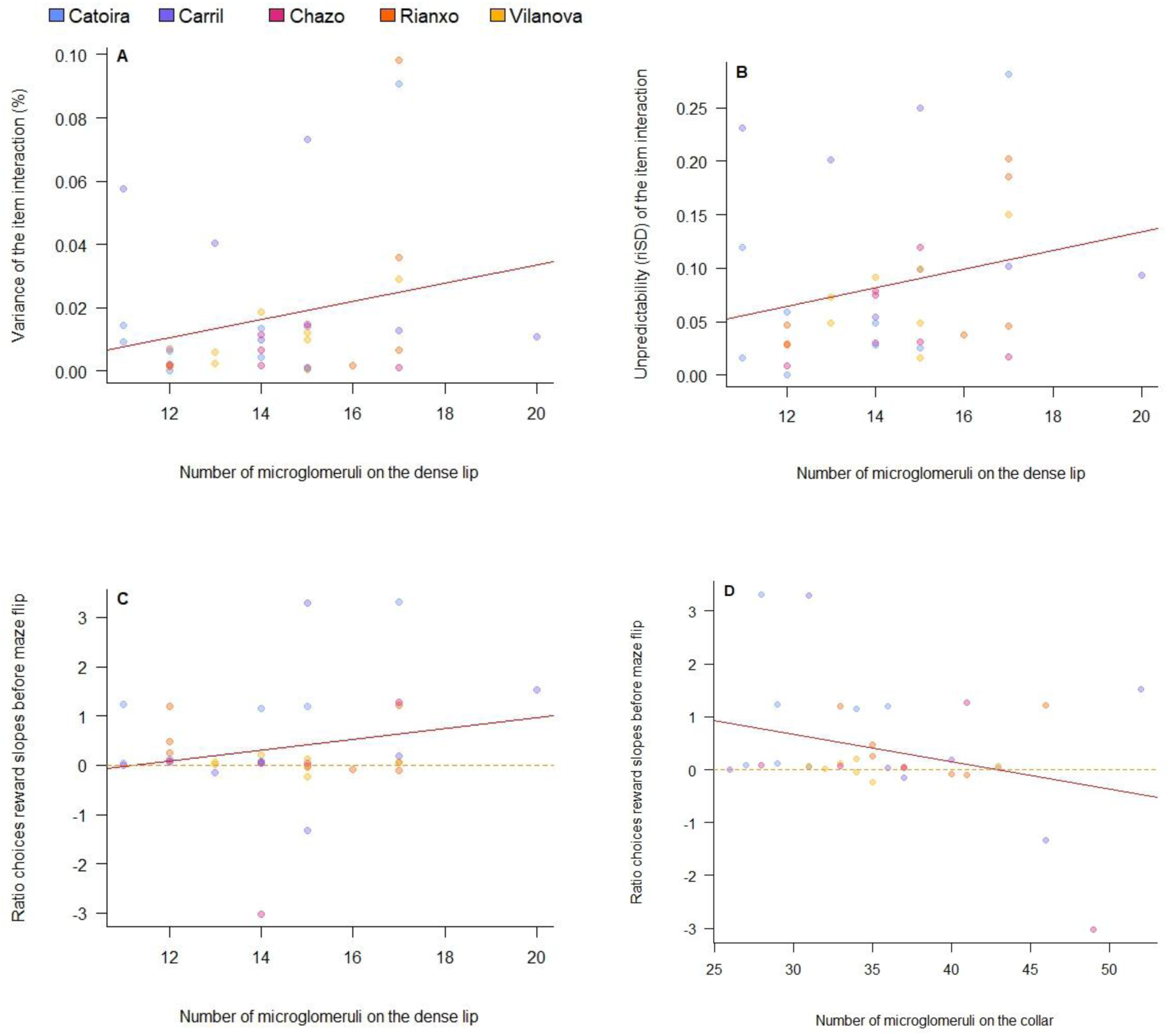
Correlation between the number of microglomeruli and behaviour of L. humile workers: correlation between the number of microglomeruli of the dense lip and the variance (**A**) and unpredictability (**B**) of the object interaction, and the slopes of choices before maze flip (**C**); correlation between the number of microglomeruli of the collar and the slopes of choices before maze flip (**D**).

## Discussion

The behavioral variability observed at the individual level suggests two mechanisms by which *Linepithema humile* colonies manage to explore new environments: through efficiency associated with body size and neural processing areas. On one hand, larger ants showed higher speed, movement, exploration and more microglomeruli on the collar (visual perception) while they did not show larger relative neuropil volumes. Smaller ants explored the arena differently, visiting its inner zone more frequently. Behavioural differences linked to body size could increase the efficiency of colony exploration due to different trial-and-error strategies. On the other hand, individuals who invested more in processing than in perception were more efficient at exploring complex environments, especially those who processed olfactory rather than visual stimuli.

Complex scenarios resolution may be favoured by this small number of individuals in whom the correlation between behaviour and neuroanatomy was observed (see dots outside the dashed line in Figure 2D-E, Figure 3C-D). This possible evolutionary strategies supports the hypothesis of the colony’s investment reduction in the behavioural and neural plasticity of its members in *L. humile* (e.g., Sanmartín-Villar et al., 2021), limiting costs to cope with complex environments through a neural demanding mechanism in only a few individuals, while in a non-neural demanding mechanism in most them. Our results suggest that exploration at the colony level might depend on the variability among-individuals, but interestingly, in the case of our study this variability was not dependent on the individual adult development - as division of labour in eusocial animals is often explained - because we analysed workers of the same age after emergence. Future studies should corroborate in the field the consequences of the mechanisms proposed here for local adaptation, invasiveness, and coping with climate change.

The expected similar behaviour among workers and lack of correlation between behaviour and neuroanatomical traits were only observed at low environmental complexity. Individuals showed different and dependent responses of their neuroanatomy when exposed to complex environmental challenges, such as novel experiences and choices. Sensory neuropils hardly affected behavior, only detecting faster exploration after object exposure in individuals with larger visual lobes which support the navigation improvement through volume increase of primary lobes as seen in other species of ants (*Cataglyphus nodus*; Grob et al., 2017, *Cataglyphis fortis*; Schmitt et al., 2016, *Camponotus rufipus*; Yilmaz et al., 2017). Processing areas (central complex and mushroom bodies) determined exploration and maze progress. Progress in the maze was positively dependent on the number of synaptic clusters in the dense lip (olfactory processing), but negatively dependent on those in the collar (visual processing). Individuals who showed more dense lip synaptic clusters showed higher behavioral variability (variance and unpredictability). These results highlight the relevance of high processing areas in the resolution of complex environments and the different functions of different brain regions. Specialized behavioral strategies could be expected from a higher neuropil volume due to the development of synaptic clusters, but also a lower number of these clusters due to pruning (the concentration of microglomeruli in a more volumetric button due to specialization; Changeux and Danchin, 1976; Cowan et al., 1984). The correlations found between exploration efficiency and the number of microglomeruli could highlight the diversity of behavioral strategies (variance, unpredictability) rather than learning processes, although they coil highlight long term memory formation due to the increase of microglomeruli density (Falibene et al., 2015).

In contrast to the expected, *L. humile* workers showed low percentage of exploratory performance, exploring an average of 60% ca. of the open field, interacting with the object up to 18% of time, and selecting rewarded paths in 27% ca. of cases. Although *L. humile* is considered a monomorphic species, body size conditioned behavioural and neural traits. The allometry for speed and body size is a common phenomenon in ants due to the higher motion of individuals with longer legs and higher metabolic rate (Hurlbert et al., 2008), but we did not expect to see such a strong difference within a monomorphic species. Assuming size-dependent differences, we expected that body size correlate with risk-taking decisions as it occurs in fish (e.g. Darby & McGhee, 2019; but see Brown & Braithwaite, 2004), turtles (Ibáñez et al., 2013), or snakes (Mayer et al., 2016). Therefore, we expected a body size dependency on the interaction with the novel object (but see the mismatch between neophobia and boldness in Carter et al., 2012), the use of the inner part of the arena (assumed as a bolder trait for some authors; Carlson & Langkilde, 2013; Detrain et al., 2019; Sneddon, 2003; Valle, 1970; Walsh & Cummins, 1976), and the participation on the maze. However, none of these traits correlated with body size, suggesting that *L. humile* workers may not evaluate the environment challenges according to their body characteristics.

The interaction with the object was less variable and unpredictable for larger workers, suggesting a more homogeneous behaviour. In addition, larger workers reached more times the reward inside the maze but decreased their ratio of choices to reward through observations. Although we could not determine the overall volume changes in the brains based on experience (see Gronenberg et al., 1996) due to our methodology, our results support the relationship between body size, brain development and foraging behaviour shown on *Atta cephalotes* (Arganda et al., 2020), highlighting our findings that larger individuals were better in simple tasks like exploration and navigation but not in complex tasks like in the maze and also being associated with larger volume ratio of primary to secondary neuropil. All these results suggest that large workers succeeded to better solve environmental complexity (higher exploration and maze-solving, more constant object interaction) due to their higher locomotive traits, but not due to the neuroanatomy that might underlie their cognitive skills (no correlation was found between speed or exploration or body size and neuropil relative volumes or microglomeruli abundance). In fact, our results suggest that large workers only had more microglomeruli in the collar than smaller workers (therefore, no more microglomeruli on the lip than smaller workers). We consider improbable that larger workers could be able to better solve the tests in which they were exposed due to their higher secondary processing of visual stimuli in comparison with the chemical stimuli (more microglomeruli on the collar, but no more on the lip; Falibene et al., 2016; Gordon et al., 2018, 2019; Gordon & Traniello, 2018; Seid & Wehner, 2008) due to the lack of visual references and similitude among choices. In any case, our results support the idea that body size can influence environmental exploration in monomorphic species as it was shown in other species such as *Lasius niger* (*e.g.*, Grześ et al., 2016; Okrutniak et al., 2020). Therefore, future behavioural studies should control for the effect of body size in monomorphic ant species but also analyse whether worker body size is related to the spatial complexity of the environment as an evolutionary strategy.

Focal workers acclimatised to the experimental setup, reducing their average speed, increasing the interaction with the object and decreasing their mobility in the maze over the observations. This suggests that ants were able to modify their behaviour according to previous experiences, as was previously shown in ants of the same species (Wagner et al., 2023). However, we cannot discard that the obtained outcomes depend on other factors such as fatigue or developmental-related changes. Previous works done with repeated observations on open fields showed different patterns (reduction of locomotion in a ten repetition test in *L. niger*, Sanmartín-Villar et al., 2021; increase of locomotion in a five repetition test in *L. humile*, Sanmartín-Villar et al., 2021). If ants recognise the object from the previous tests, the object introduced could only be considered as novel on the first observation. Despite the lack of novelty and apparently lack of functionality of the object (except for climbing), the focal workers interacted more with it through the observations, especially the workers that showed higher exploration, while the opposite happened in the maze, in which workers could find a reward. Note that we did not starve focal individuals, and a lack of food motivation could interfere with performance in the maze. Still, we expected that ants would increase their maze performance while losing the fear to navigate through it and especially after the reduction of their speed through observations, which may could be due to the speed-accuracy trade-off and its dependency on neural structures (reviewed in Standage et al., 2014).

The increase of the environmental complexity through the addition of an object in a relatively known arena (experienced through 10 minutes) triggered behavioural differences among workers and modified the apparently characteristic homogeneous locomotion pattern of the *L. humile* workers in open fields (also seen in Sanmartín-Villar et al., 2021). The unpredictability of the object interaction, *i.e.*, its deviation through the expected pattern, correlated with the neural area in charge of processing chemical stimuli (dense lip) and the mean and variance of the object interaction correlated with the relative volume of the central complex, which is in charge of navigation (Honkanen et al., 2019; reviewed in Webb & Wystrach, 2016). The increase of the object interaction and movement (marginally exploration) could fit with the expected result of environmental enrichment (reviewed in Newberry, 1995), in which individuals improve their cognitive skills after the addition of objects. Individuals exposed to the object used more the inner zone of the area in posterior tests when the object was not present while control individuals showed this pattern with a delay of 10 min, suggesting an improvement of the area exploration after previous stimuli. However, we cannot discard that the behaviours observed were caused by the stress produced when adding a new element in a known area. Considering the correlation trend of the object interaction with the central complex (navigation centre), we propose a third explanation in which workers used the object as a spatial reference to which they recurred to orient in an arena lacking other spatial clues.

The difficulty of the maze test highlighted the relation between neuroanatomy and behaviour. Within the mushroom bodies (in charge of information processing, memory and learning), the maze performance before the flip was only correlated with the relative volume of the medial calyx, and not with the lateral calyx. As medial and lateral calyces are considered to perform the same function, our results might be a suggestion for further research trying to disentangle distinctive information processing among them. The relative volume of the central complex (navigation centre) also correlates with maze performance before the flip. The difficulty of the maze due to its design and the lack of references accomplished the aim of exposing ants towards an exercise focused on exploration rather than learning (we did not train the ants before the tests). Workers with more microglomeruli in dense lip showed higher ratio of choices to reward before maze flip, which suggests that the presence of more synaptic routes - but smaller - in this region responsible for chemical processing allowed individuals to improve their efficiency on the maze. We discarded that the experience on the maze was responsible for the synaptic cluster development during the tests because of the similar microglomeruli found between control and focal workers, although more research must be performed to conclude it. In contrast, the number of microglomeruli of the collar negatively correlated with the ratio of choices to reward before maze flip. In addition, the relative volume of the central complex (more relevant in the maze performance) was negatively correlated with the relative volume of the optic lobes. We interpret these results as a conflict between the role of primary and higher order processing regions in the studied behaviours: one region in charge of perceiving and processing of visual stimuli (optical lobes), and two in charge of the higher order processing of stimuli (the chemical information in the non-dense lip in the mushroom body calyces, and the visual, chemical and mechanosensory information required for orientation in the central complex). Individuals that relied more on visual stimuli were probably not able to disentangle the orientation on the maze due to the exact symmetry of all the choices. However, individuals relying more on the chemical stimulus could better interpret their own pheromones or other odors present in the maze (although backwards behaviour was uncommon). Surprisingly, no correlations were found with the antennal lobes, which we expected to have a higher relevance in the ants’ orientation.

The relevance of the microglomeruli of the dense lip in behaviour (unpredictability of the object interaction, increase of rewarded path choice) opposes the results found in honey bees (Van Nest et al., 2017), in which only the microglomeruli of the non-dense lip (also olfactory) were related with foraging behaviour. Thermal fluctuations affect the microglomeruli density in the non-dense lip in the mushroom body calyx of *Camponotus mus* but were less evident in dense lip and visual collar (Falibene et al., 2016). If the correlations between these experiments are comparable, this would highlight phylogenetic differences in the neuroethology of the species, with behaviour being dependent on different neural structures across insect and ant species. However, all these experiments highlight the expected relevance of the olfactory stimulus over the visual one to drive behaviour.

Our results suggest that individual variability depends on neuronal development and highlight behavioural plasticity and the strategies to cope with environments with different degree of challenge in *Linepithema humile*, skills likely related with its adaptation in introduced areas. Although we could not observe the associated neuronal changes with each exposure to behavioural tests, likely to decipher in detail, the mechanisms involved in their behavioural patterns, we provide the first neuroethological clue to understand the invasive strategies of this species. We attempted to dissect and correlate the behavioural and neuronal plasticity of an invasive species to cope with novel scenarios, providing a framework for understanding the dynamics for colony fitness in the introduced areas. We are convinced that the present results could be the basis for future control and management programs dealing with *L. humile* invasion. For instance, neural disruptors could affect the behaviour of worker ants differently depending on their size and on the complexity of their environment.

## Acknowledgements

We are grateful to Xoan Sanmartín Pazos for his help in the field work and to the staff of the Łomna Biological Station (MiIZ) for allowing us to produce uninterrupted work even in national holidays and heavy snow conditions. This research is part of the project No. 2021/43/P/NZ8/03306 co-funded by the National Science Centre and the European Union Framework Programme for Research and Innovation Horizon 2020 under the Marie Skłodowska-Curie grant agreement no.945339. Part of this research was supported by the 2017 program for attracting and retaining talent of Comunidad de Madrid (2021-5A/BMD-20960). For the purpose of Open Access, the author has applied a CC-BY public copyright licence to any Author Accepted Manuscript (AAM) version arising from this submission.

## Contributions

M.W. and I.S.V. conceived the ideas and designed the methodology; V.C and I.S.V. collected the samples; S.N, V.C., and I.S.V conducted the experiments; M.E.V. and S.A. performed the protocols and conducted the brain imaging; S.N collected the behavioural data; M.E.V., I.A.C., and S.A. collected the brain-related data; I.S.V. analysed the data; S.N., M.E.V., S.A., I.A.C., and I.S.V. wrote the manuscript; M.W., S.A., and I.S.V. lead the funding acquisition, project administration, and supervision. All authors contributed critically to the drafts and gave final approval for publication.

## Notes

### Competing Interest Statement

The authors have declared no competing interest.

### Summary of Updates

Authors may wish to note any significant differences between this version and previous versions of the manuscript. Author affliations were updated accordingly.

