## Supplementary material for "Complexity shapes uniqueness: Neuropil volumes and synaptic clusters shape behavioural plasticity under challenging environments in the invasive Argentine ants" (from another version of this preprint: version 1, 2025-03-13)

*Maze performance*

Ants tended to participate in the maze. Ants left the starting point to explore the maze in 70.90% of the 850 observations made. Most of the ants (83.76%) entered the maze between three to five observations. When the ants decided not to participate in the maze, it was mainly on the first observation (participation in each observation: 1= 68.63%, 2= 77.45%, 3= 84.31%, 4= 79.41%, 5= 90.20%; $\chi^2_4$= 9.37, *p*= 0.052). However, the participation of ants across observations was not statistically different ($\chi^2_4$= 4.00, *p*= 0.406). The ant size did not affect the ant participation in the maze ($F_{1;149}$= 191, *p*= 0.169).

Ants' path selection was conditioned more by the type of path (open, closed) than by the side (left, right) or previous experiences, except when the maze was flipped. Ants choose more the rewarded path when starting the maze for the first time ($\chi^2_1$= 8.76, *p*= 0.003; N= 148). The first path selected the first time that ants participated in the maze was not influenced by innate lateralization (left path: 47.94%, right path: 52.03%; N= 148, $\chi^2_1$= 0.24, *p*= 0.622) nor conditioned by the first direction selected in the previous observation in the second, third and fifth observations (same: 50.73±0.37%; different: 49.27±0.37%). However, when the maze was flipped (fourth observation), 65.85% of focal ants chose the same first path as they did in the previous observations (N= 54; $\chi^2_1$= 8.24, *p*= 0.004; Suppl. Figure 1A).

A similar number of ants increased or decreased the ratio of choices to reward through observations both before (positive slopes: 55.77%, N= 87; negative slopes: 44.23%, N= 69) and after (positive slopes: 61.90%, N= 91; negative slopes: 44.22%, N= 65) flipping the direction of the maze. After the maze flip, half of the individuals maintained their strategy performed before the maze flip, increasing (25.00%, N= 39) or decreasing (27.56%, N= 43) the ratio of choices to reward; while the other half changed their pattern, increasing before and decreasing after (30.77%, N= 48) or decreasing before and increasing after (16.67%, N= 26; Suppl. Figure 1B). Ants that increased the ratio of choices to reward before the maze flip equally selected to continue or to change their strategy after the maze flip ($\chi^2_1$= 0.93, *p*= 0.335). However, almost twice as many individuals continued to decrease the selection of rewarded choices after the maze flip compared to the number of ants that increased it ($\chi^2_1$= 4.19, *p*= 0.041). Ants participating in the maze returned 5.71±3.67 times to one of the paths already visited. This value was constant across observations (no intraindividual variability; Z= -1.32, *p*= 0.188), showed no differences among individuals (interindividual differences; ΔAIC= -4) and no differences according to the orientation of the maze ($\chi^2_1$= 0.00, *p*= 0.986).

More than half of the focal individuals (63.46%) reached the end of the maze at least once. The ants rarely reached the end of the maze more than twice (number of individuals according to the number of times they reached the maze end: $N_0$= 57, $N_1$= 43, $N_2$= 33, $N_3$= 12, $N_4$= 8, $N_5$= 3; S= 70, *p* = 0.003). Half of the ants needed more time to reach the end of the maze after reaching it once (44.64%, N= 25) while half needed less time (55.36%, N= 31). Ants that never reached the end of the maze or did it once were smaller than those that did it more than once ($\chi^2_2$= 6.34, *p*= 0.012) and to those that decreased the ratio of choices to reward through observations (Z= 2.65, *p*= 0.024; Suppl. Figure 2C).

The design of the maze used was probably difficult to solve for the ants. Although workers were able to detect the paths allowing maze exploration and almost never came back from the visited paths, the success in reaching the end was very low, with only 2% of workers finding the reward through all observations and only half of the focal ants improving the time to reach the reward after finding it in previous observation. In general, workers showed no influence on the choices selected in previous observations and no specific strategy on selecting paths. However, the flip of the maze disturbed workers who reduced the choices allowing maze exploration, forcing most of them to worsen their performance.

The design of the maze evidenced that the focal workers did not have a lateralized preference for dichotomous choices, either in their first selection or a fixed lateralized pattern of choices (if this was the case, ants that chose to go always to one side would quickly reach the end of the maze) and changed their choice throughout their exploration.

*Repeatability*

The exposure to complex environments disrupted the exploration repeatability of ant workers and made them repeatable in the use of added objects. Open field exploration was almost three times more repeatable in control ants (R= 0.65±0.16, CI [0.20, 0.84]) than in focal ants (R= 0.22±0.04, CI [0.14, 0.30]). In contrast, control ants showed no repeatability in the use of the inner area of the arena in the object exposure tests (no object; R= 0±0.08, CI [0, 0.27]) while focal ants showed similar repeatability when interacted with the new object than the obtained for exploration in open field (R= 0.24±0.04, CI [0.17, 0.32]). Ants showed a non repeatable ratio of choices to reward (R= 0.02±0.02, CI [0, 0.07]).

Although this work was not aimed to unravel differences on the use of coefficients of variation and formulas based on the ICC to calculate repeatability but without considering measurement errors (Dingemanse et al., 2022), we point out the discrepancy among both methods using the obtained outcomes. Both methods differentiated the variability of control and focal workers' using the border and the inner area of the arena, but exploration was three times more repeatable in control than focal workers (but with CI slightly overlapped) while its coefficients of variation were similar. Both methods produced similar outcomes when the environment differed (with and without object) but outcomes differed (although statistically) when the environment was similar (open field). We assume that the differences found using repeatability could be linked to uncontrolled measurement errors as suggested by the authors of its critics (Dingemanse *et al.* 2022). Although control workers were randomly tested within the focals and following similar procedures, they were not handled among tests nor exposed to new objects (object and maze), which could influence the differences found in measuring repeatability.

**Supplementary Figure 1**. Maze performance. **A**: Number of focal ants that selected the same (orange) or different (blue) first choice as in the previous test. **B**: Strategies performed during the Maze tests according to the relative number of choices allowing maze exploration. Increase choices (orange triangles): number of individuals that increased the relative number of choices to reward through observations (three observations before maze flip, two observations after). Decrease choices (blue triangles): number of individuals that decreased the relative number of choices to reward through observations (three observations before maze flip, two observations after). The width of the bars represents the percentage of individuals who continue or change strategy before and after the maze turn. *: significant differences among strategy selection. **C**: Body size of individuals reaching the end of the maze in different times (less than two, more than one) considering different kinds of performance (worsen or improve the time needed to reach the end in a posterior observation).

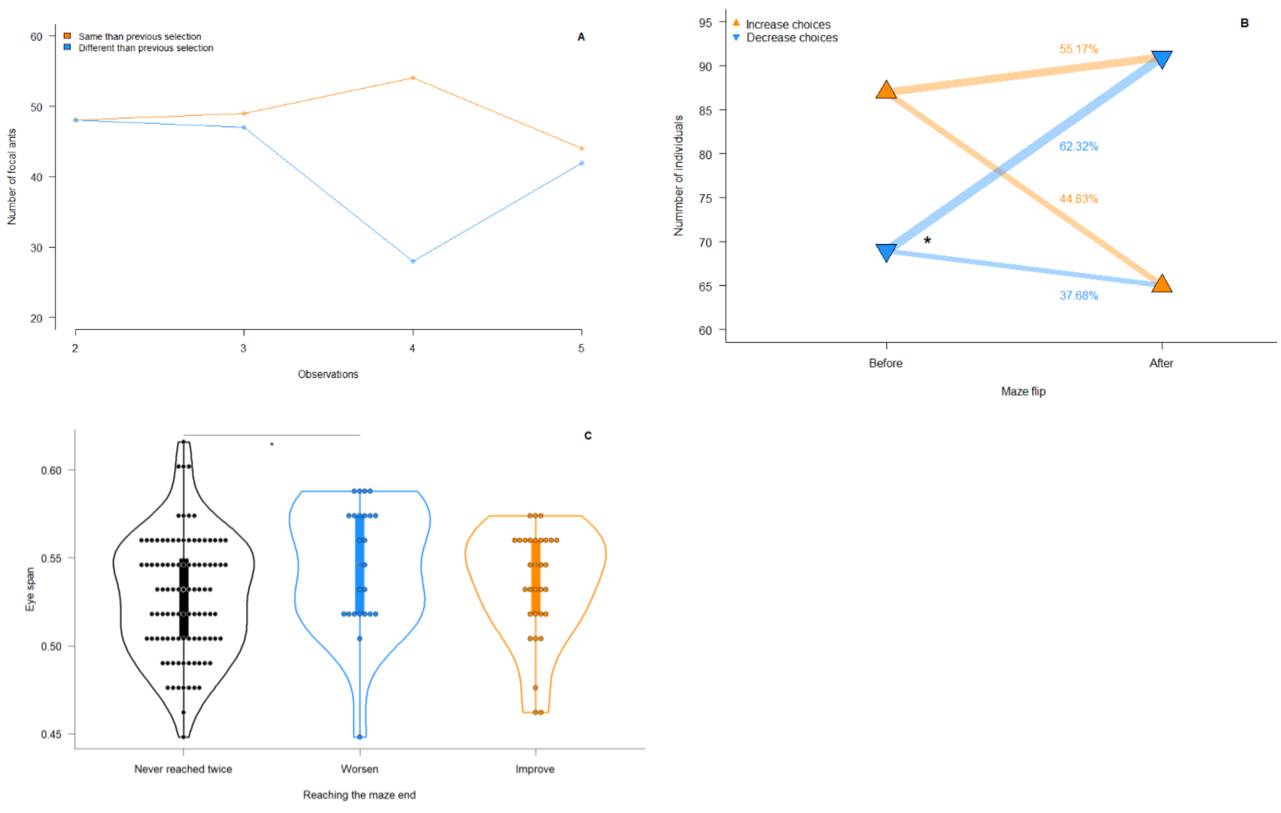
